# Semantic object-scene inconsistencies affect eye movements, but not in the way predicted by contextualized meaning maps

**DOI:** 10.1101/2021.05.03.442533

**Authors:** Marek A. Pedziwiatr, Matthias Kümmerer, Thomas S.A. Wallis, Matthias Bethge, Christoph Teufel

## Abstract

Semantic information is important in eye-movement control. An important semantic influence on gaze guidance relates to object-scene relationships: objects that are semantically inconsistent with the scene attract more fixations than consistent objects. One interpretation of this effect is that fixations are driven towards inconsistent objects because they are semantically more informative. We tested this explanation using contextualized meaning maps, a method that is based on crowd-sourced ratings to quantify the spatial distribution of context-sensitive ‘meaning’ in images. In Experiment 1, we compared gaze data and contextualized meaning maps for images, in which objects-scene consistency was manipulated. Observers fixated more on inconsistent vs. consistent objects. However, contextualized meaning maps did not assigned higher meaning to image regions that contained semantic inconsistencies. In Experiment 2, a large number of raters evaluated the meaningfulness of a set of carefully selected image-regions. The results suggest that the same scene locations were experienced as slightly *less* meaningful when they contained inconsistent compared to consistent objects. In summary, we demonstrated that – in the context of our rating task – semantically inconsistent objects are experienced as less meaningful than their consistent counterparts, and that contextualized meaning maps do not capture prototypical influences of image meaning on gaze guidance.

## Introduction

Visual processing varies as a function of the retinal location at which a stimulus is presented: with increasing eccentricity, processing is affected by crowding and a decrease in resolution (see Rosenholtz, 2016 and Stewart et al., 2020 for reviews). Being able to rapidly move the central parts of the eyes is therefore necessary to extract fine detail across large parts of the visual field. Consequently, eye movements are critical for visual processing and it is important to understand what processes underpin gaze guidance. Currently, the most popular framework for answering this question assumes that the factors influencing human gaze allocation belong to two broad categories: (i) image-computable features of the input processed in a bottom-up fashion, and (ii) the internal states of the individual, such as knowledge or intentions, exerting their influence in a top-down manner (Berga & Otazu, 2020; Henderson & Hayes, 2017; Kollmorgen et al., 2010; Rothkopf et al., 2016).

Support for the notion that image-computable aspects of the input are important for the guidance of eye movements comes from studies demonstrating that where humans look in images can often be predicted by analyzing the visual features of these images (Borji et al., 2013). Algorithms generating such predictions are called saliency models. Early saliency models, such as GBVS (Harel et al., 2007), AWS (Garcia-Diaz, Fdez-Vidal, et al., 2012; Garcia-Diaz, Leboran, et al., 2012) or the model by Itti and Koch (Itti & Koch, 2000; see also Krasovskaya & MacInnes, 2019), attempted to maximize the accuracy of their predictions relying on simple features such as intensity, color, and orientation contrasts. While the predictive power of these models was moderate (Kümmerer et al., 2015), state-of-the-art saliency models, based on powerful machine-learning algorithms called deep neural networks (see Storrs & Kriegeskorte, 2019 for review), can predict fixation locations much better than their predecessors while still relying exclusively on image features (Kümmerer et al., 2017). One fundamental difference is that while earlier models were based on parameter values determined by hand, current models such as DeepGaze II (Kümmerer et al., 2016, 2017) or MSI-Net (Kroner et al., 2020) are based on supervised learning, which does not require explicitly defined parameter values.

One limitation of all saliency-based approaches is their difficulty to account for factors in oculomotor control that are not image-computable (Bayat et al., 2018; Bruce et al., 2015; Henderson & Hayes, 2017; Pedziwiatr et al., 2021a; Tatler et al., 2011). For example, the fixation-patterns of individuals viewing the same stimulus can vary as a function of their task and goals (Hoppe & Rothkopf, 2019; Koehler et al., 2014; Rothkopf et al., 2016; Yarbus, 1967). Importantly, however, oculomotor behavior is not constantly subjugated to a task; humans (and many other animals) are intrinsically motivated to obtain information, and often move their eyes with no purpose other than to explore the environment (Gottlieb & Oudeyer, 2018). Both early (Itti & Koch, 2001) and more recent work (Adeli et al., 2017; Veale et al., 2017; Zelinsky & Bisley, 2015) argues that the oculomotor behavior exhibited in such ‘free-viewing’ conditions can be largely explained by image-computable features.

This contention has not remained unchallenged. A number of studies demonstrated that even when observers view images without a task, the spatial allocation of fixations can be guided by factors which are not captured by current saliency models, namely, the semantic content of the visual scene (Henderson et al., 2019; Peacock et al., 2019; Wu et al., 2014). One well-studied semantic effect in eye movement research relates to object-scene consistency, where eye movement behavior changes depending on the extent to which objects are semantically consistent with the scene. In a seminal study (Loftus & Mackworth, 1978), one example stimulus showed a farmyard scene either with a (semantically consistent) tractor, or a (semantically inconsistent) octopus. Inconsistent objects such as the octopus were looked at earlier, attracted more fixations, and were inspected for longer in comparison to consistent objects. While some mixed results have since been found with respect to the timing of eye movements (Wu et al., 2014), there is robust evidence demonstrating that object-scene inconsistencies lead to more and longer fixations (Coco et al., 2020; Friedman, 1979; Henderson et al., 1999; Öhlschläger & Võ, 2017; Pedziwiatr et al., 2021a).

Two primary mechanisms have been proposed to explain these effects. First, objects that are viewed in inconsistent contexts are processed less effectively, as indicated by the drop in recognition (Munneke et al., 2013) and detection (Biederman et al., 1982) performance (see also Kaiser et al., 2019). Consequently, more fixations towards, and longer inspection times of inconsistent objects are thought to reflect the increased resources needed to process these stimuli (Bonitz & Gordon, 2008; Friedman, 1979). A second, and not mutually-exclusive, explanation for the effects of object-scene inconsistencies on eye movements is based on the notion that inconsistent objects are *“more informative”* (Loftus & Mackworth, 1978), *“semantically informative”* (Henderson, 2011; Henderson et al., 1999), or *“contain greater meaning”* (Peacock et al., 2019). According to this idea, people look at inconsistent objects in an effort to maximize extraction of meaning from a scene.

This second interpretation has recently gained increased attention, in particular with the development of meaning maps, a method to quantify the spatial distribution of ‘meaning’ across an image (Henderson & Hayes, 2017, 2018). Meaning maps are created by first partitioning an image into many circular, partially-overlapping patches. These patches are presented to individuals, who view them without knowing the scene from which they were extracted (hence these maps are called context-free). Participants are asked to use a Likert scale to *“assess how “meaningful” an image is based on how informative or recognizable”* they think it is. Finally, these ratings are combined into a smooth distribution over the image to create a map. Meaning indexed by this method has been demonstrated to be a better predictor of fixations than a simple saliency model. This finding has been interpreted as evidence that semantic information rather than image-computable features control eye movements (Henderson & Hayes, 2017, 2018). The meaning map approach is rapidly gaining popularity, and has been used to study eye movements in various contexts (listed in Henderson et al., 2021).

A recent study evaluating the meaning map approach and comparing them to a wider range of saliency models highlights some limitations of the method (Pedziwiatr et al., 2021a; see Henderson et al., 2021 and Pedziwiatr et al., 2021b for ongoing debate). First, the findings demonstrate that meaning maps are outperformed in predicting fixations by DeepGaze II (Kümmerer et al., 2016, 2017), a saliency model based on a deep neural network, that indexes high-level features rather than meaning. Second, it was found that meaning maps in their original form do not ascribe more meaning to scene regions occupied by objects that are semantically inconsistent with the global scene context compared to consistent objects presented in the same region and matched in terms of low-level features. Together, the results of this study led to the conclusion that there is so far no evidence that meaning maps measure semantic information *per se* (for further discussion see Pedziwiatr et al., 2021b). Rather, they might index visual features that can be correlated with semantics. In this respect, the original form of meaning maps are similar to modern saliency models.

As detailed above, the original meaning maps ignore the global context of the scene – they are created from ratings of isolated, ‘context-free’ image patches. To resolve this issue, Peacock et al. (2019) recently proposed *contextualized* meaning maps to allow meaningfulness ratings to capture global scene context effects, such as object-scene inconsistencies. Contextualized meaning maps differ from the original meaning maps in one important detail: during rating, each patch is presented alongside the full scene from which it originated. Therefore, raters have access to the global scene context when assessing the meaningfulness of the patch. Given the critical importance of context in scene semantics (Biederman et al., 1982; Võ et al., 2019), contextualized meaning maps might be better suited to quantify semantic information within visual scenes. Surprisingly, Peacock et al. (2019) found that contextualized meaning maps predicted gaze density in a free-viewing task equally well as context-free meaning maps (and both predicted gaze density better than the GBVS saliency model). They suggested, however, that dissociations in prediction performance between context-free and contextualized meaning maps might only occur for scenes containing object-scene inconsistencies.

In the current study, we therefore assessed the extent to which contextualized meaning maps are sensitive to semantic object-scene inconsistencies. Specifically, if inconsistent objects are more meaningful (Henderson, 2011; Henderson et al., 1999; Loftus & Mackworth, 1978; Peacock et al., 2019), then contextualized meaning maps should assign higher meaning to regions occupied by them, and this should predict increased fixations on these objects (relative to consistent objects). Using exactly the same procedure and instructions as Peacock and colleagues (2019), we created contextualized meaning maps for two types of indoor scenes, which were identical except for one object (Öhlschläger & Võ, 2017). This object was either semantically consistent with the context, such as a hair brush on a bathroom sink, or the object was replaced with an inconsistent object, such as a shoe on the sink. We conducted a detailed analysis of these maps across scene types, and compared them to fixation patterns of human observers.

To anticipate our findings, we demonstrate that contextualized meaning maps are not able to predict the gaze changes elicited by the manipulation of semantic object-context consistency. Moreover, our first experiment provided initial evidence that contextualized meaning maps might attribute *less* meaning to scene regions that contain inconsistent compared to consistent objects. Given this surprising result, in a second experiment, we asked a large number of raters to provide meaningfulness ratings for a carefully controlled set of image patches. The results of this second experiment replicated the surprising result from the first experiment, showing that semantically inconsistent objects are judged as slightly less meaningful than consistent objects. Overall, these results call for the assumptions of the meaning map approach to be reconsidered.

## Methods and Results

### Experiment 1

The main goal of Experiment 1 was to assess the extent to which contextualized meaning maps and human fixations are sensitive to local changes in semantic information within a scene, resulting from the presence of objects that are semantically consistent vs. inconsistent with the overall scene-context. This experiment compares contextualized meaning maps to the data collected in (Pedziwiatr et al., 2021a); therefore, more methodological details on the stimuli and eye movement data can be found in that report.

#### Stimuli

The stimulus set consisted of photographs of 36 indoor scenes, taken from the SCEGRAM dataset (Öhlschläger & Võ, 2017). Each scene was photographed in two conditions: Consistent and Inconsistent, resulting in two images per scene (72 images in total). Images from the Consistent condition contained only objects that are typical for a given scene context. In the Inconsistent condition, one of these objects was replaced with an object unusual in the context provided by the whole scene, thus introducing a semantic inconsistency. For example, in one of the scenes, a hair brush on a bathroom sink (Consistent condition) was replaced with a flip-flop (Inconsistent condition) – see Fig. 1A. The SCEGRAM dataset is constructed in such a way that, across scenes, consistent and inconsistent objects are matched for low-level properties (Öhlschläger & Võ, 2017). In each scene, consistent and inconsistent objects occupy the same image locations, and the superposition of the bounding boxes of both conditions constituted what we call a Critical Region. These Critical Regions are important for the data analyses we report further below because they contain the only image regions that differ between conditions.

**Fig. 1.**
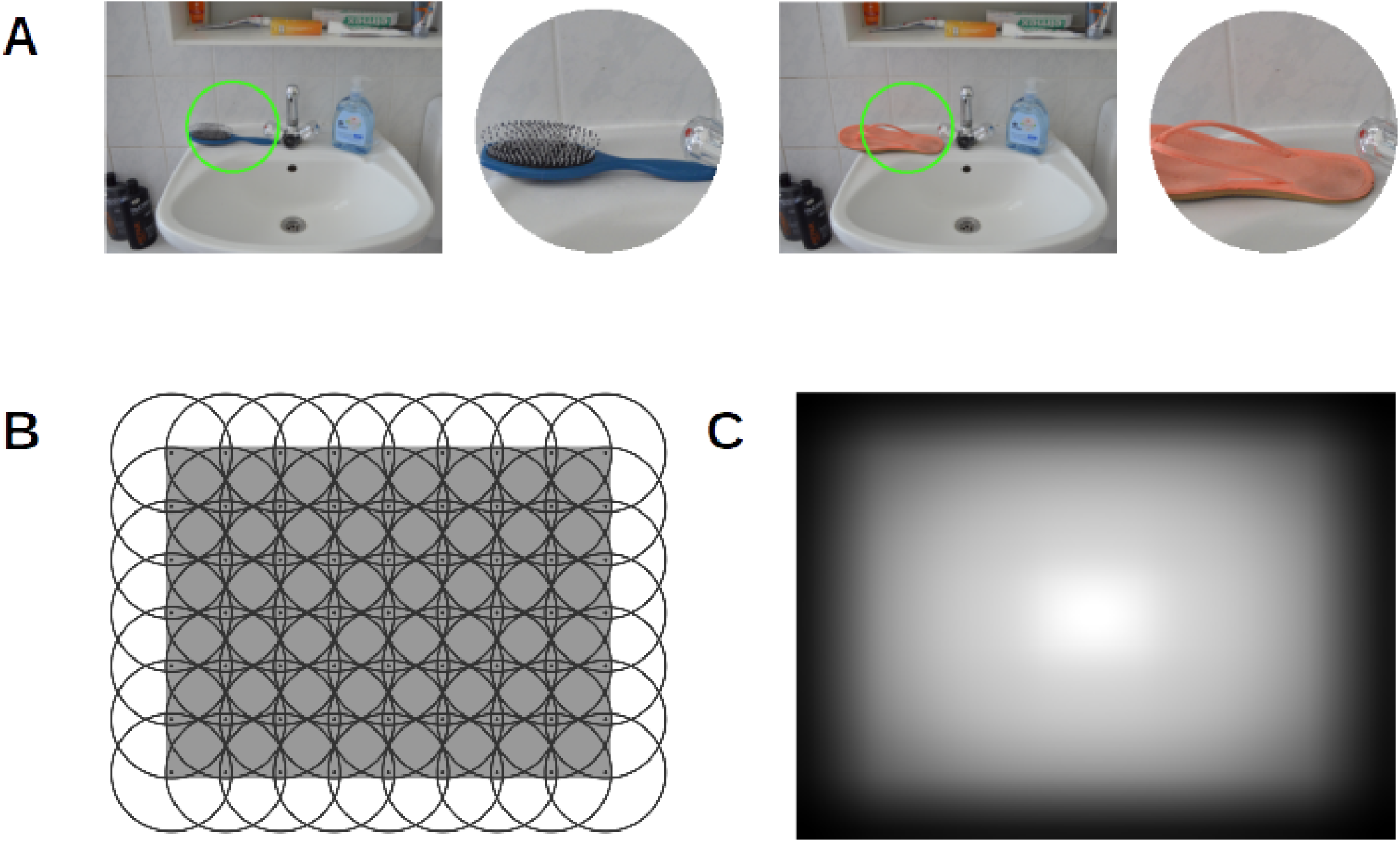
*Generating contextualized meaning maps. A) Sample stimuli from the patch-rating task used for creating contextualized meaning maps. The patch, which raters were asked to rate for its meaningfulness, was always presented next to the image from which it originated to provide the relevant context. A green circle on the context image indicated the location of the patch. Both panels show the same scene, photographed in the Consistent (left part of the panel) and in the Inconsistent (right part) condition. The images on both panels differ only with respect to the object shown in the patch. The hair brush on the left part is a semantically consistent object for a bathroom scene, the shoe on the right part is semantically inconsistent. In the task, raters were asked to assess the meaningfulness of the patches based on their informativeness and recognizability by means of selecting a value on a six-point rating scale. B) Grid used to segment images into coarse patches. Grey rectangle represents image area. C) Center bias model used in contextualized meaning maps. To account for the human tendency to allocate fixations predominantly to central image-regions (a so-called center bias), contextualized meaning maps assign different weights to different pixels of the maps depending on their location. This re-weighting is done by computing a pixel-wise product between the maps and a model of center bias shown on this panel, in which brighter pixels indicate higher pixel-weights. See* Creating contextualized meaning maps – modeling center-bias section *for details*.

#### Eye-movement data

For all 72 images, we collected eye-tracking data from a group of 20 observers. Each observer free-viewed the full set of images displayed in a random order while their eyes were tracked with an EyeLink 1000+ eye-tracker. The images had a width of 688 pixels and a height of 524, corresponding to, respectively, 19.7 and 15 degrees of a visual angle. Each image was presented for 7 seconds, which is similar to the presentation duration of 8 s used in the original contextualized meaning maps study (Peacock et al., 2019).

To analyze the eye-movement data, fixation locations were extracted from raw eye-tracker recordings using a standard EyeLink algorithm. The discrete fixations on each image were transformed into continuous distributions by means of Gaussian smoothing (filter cut-off frequency: −6 dB; implemented in Matlab – see Kümmerer et al., 2020) followed by a normalization to the [0-1] range.

#### Creating contextualized meaning maps – overview

The procedure of creating contextualized meaning maps is identical to that used to generate the original meaning maps except that raters see the entire original image alongside the patch that they are asked to rate. We closely followed the procedure described in detail in previous publications (Henderson & Hayes, 2017, 2018; Peacock et al., 2019; Pedziwiatr et al., 2021). In summary, a pre-defined grid is used to segment the image into circular, partially overlapping patches (Fig. 1B). Next, in a crowd-sourced online experiment, each patch is presented next to the image from which it was derived, and human raters are asked to rate the meaningfulness of the patch. Presenting the full image next to the patch ensures that the rater has access to the scene context when providing their responses (Fig. 1A; see this figure for details of the rating procedure itself). Each individual patch is rated by three individuals. In our study, we used the same instructions for raters as the original contextualized meaning maps study (retrieved from https://osf.io/654uh). Specifically, human raters were asked to rate how ‘meaningful’ a patch is on a six-point Likert scale given how “*informative or recognizable*” they find it (see caption for panel A on Fig. 1 for details). To provide raters with anchoring points for their ratings, they viewed examples of patches during the instructions that should be rated as low or high (again, the same as in the study by Peacock et al., 2019). After data collection, the ratings from individual patches are combined into a smooth distribution over the image by means of averaging and interpolation. For each image, these three steps are conducted twice: once for bigger ‘coarse’ patches and once for smaller ‘fine’ patches. The maps resulting from coarse and fine patches are averaged. Finally, the regions of the average map close to the edges of the image are down-weighted (Fig. 1C). This manipulation accounts for the center-bias of human eye-movements, i.e., the tendency to look more at the central region of an image (Tatler, 2007).

#### Creating contextualized meaning maps – parameter value selection

When creating contextualized meaning maps for our stimuli, the aim was to match as closely as possible the procedure used in the original study by Peacock and colleagues (2019). Our images, however, differed in size from the stimuli used in that study and were viewed from a different distance during the eye-movement data collection. In order to account for these differences, we matched the two studies with respect to the size of coarse and fine patches in degrees of visual angle (deg), and with respect to patch density of coarse and fine patches expressed in the number of patches per square degree of visual angle (p/deg^2^). Under the constraint that the centers of each two adjacent patches have to be equidistant horizontally and vertically, these four values fully specify the grids necessary for creating contextualized meaning maps. In terms of absolute values, matching the two studies with respect to these parameters was perfect for patch diameter and resulted in 5.26 deg (coarse patches) and 2.26 deg (fine patches), which corresponded to 187 pixels and 79 pixels, respectively (205 and 87 pixels in the original study). The patch densities closest to the original we could possibly achieve were 0.56 p/deg^2^ and 0.21 p/deg^2^ (compared to 0.57 p/deg^2^ and 0.2 p/deg^2^ in the original study). Given the size of our stimuli, these values correspond to 63 coarse and 165 fine patches per image. The resulting grid for creating coarse patches if shown on Fig. 1B.

#### Creating contextualized meaning maps – data collection

The procedure described in the previous sections resulted in a total of 16 416 patches (4 536 coarse and 11 880 fine patches). As described in more detail above and in the caption for Fig. 1, each patch was rated for its meaningfulness by three human raters on a six-point Likert scale. Patches were divided into 54 sets of 304 patches each, and each set was assigned to three different raters (see details below).

Recall that each scene was photographed in a Consistent and an Inconsistent version, differing only with respect to the identity of a single object. If the raters were to view the same scene in both versions, there would be a high chance that they might guess the main focus of the study and, in turn, adjust their rating strategy (by, for example, conditioning all rating values on the presence – or absence – of the semantic inconsistency in the context image). To ensure that meaning maps in scene pairs were independent, we assigned patches to sets in such a way that each rater never saw the same scene in both the Consistent and Inconsistent conditions. Specifically, we divided all the patches into two subsets. The first contained half of the patches from the Consistent condition and half from the Inconsistent, with the patches in both these halves derived from different scenes. The other subset contained the remaining patches. Patches in each set presented for rating were always drawn only from one of these subsets. Within each subset, patches were allocated to the Consistent and Inconsistent condition randomly. Because of this division, raters were never exposed to the same scene in both conditions but each rater was still exposed to scenes with and without semantic inconsistencies.

Each set was rated by three unique raters, and 162 raters were recruited in total. The order of patch presentation was randomized for each rater separately. Data collection was conducted online. The raters were recruited using the crowdsourcing platform Prolific (www.prolific.co) and the patch-rating task was implemented as a Qualtrics survey (Qualtrics, Provo, UT). All our raters had to meet the following eligibility criteria: they had to be of U.S. nationality (as in the original contextualized meaning maps study), they had to have submitted at least 100 tasks to Prolific before, had to have an approval rate of 95% or more, and had to use a laptop or a personal computer to complete the task. They were financially reimbursed for their time and were allowed to participate in our study only once. Median completion time was 17.08 minutes (interquartile range: 9.19).

#### Creating contextualized meaning maps – modeling center-bias

Recall that the final step of creating contextualized meaning maps involves reweighting the map with a model of center bias. Such models have the form of smooths distributions over the image, with higher values closer to the image center (Clarke & Tatler, 2014). When creating contextualized meaning maps, we followed the original authors and relied on a model provided with the saliency model GBVS (Harel et al., 2007; to be precise, we used the inverse of the center-bias model included in the *invCenterBias.mat* file; inversion was achieved by subtracting all values from one). This model is shown on Fig. 1C, its effects are illustrated on Fig. 2D and E.

**Fig. 2.**
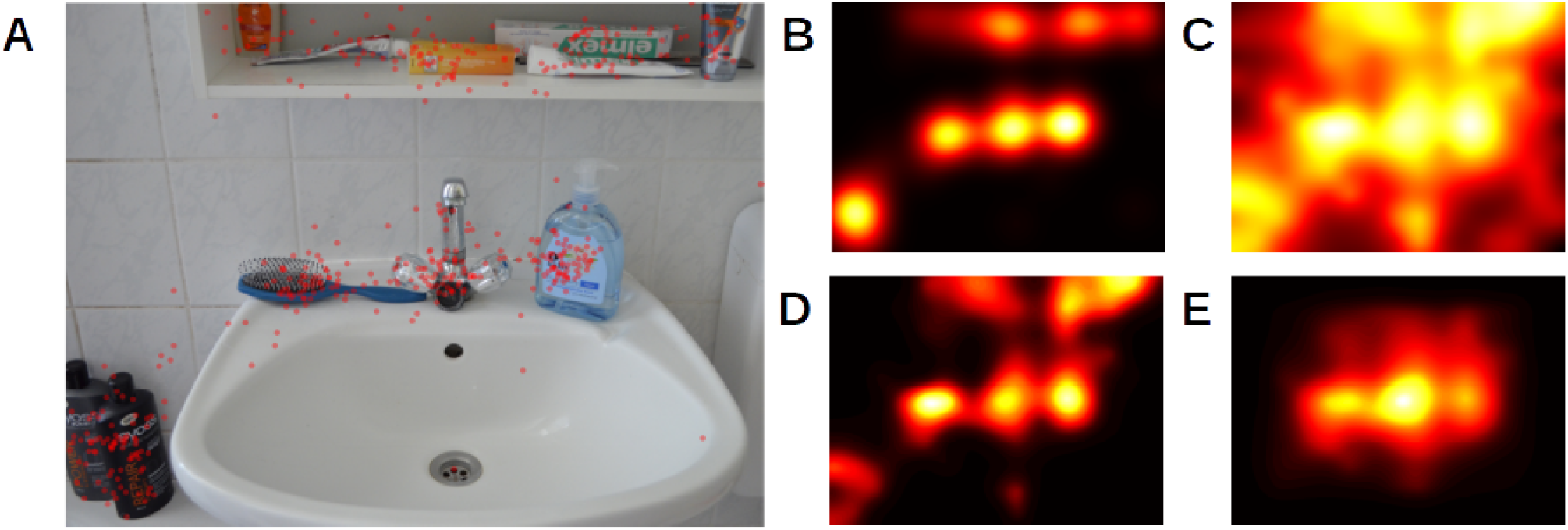
*Gaze data and outcomes of selected steps of creating a contextualized meaning map for an example scene. A) Singles scene from the Consistent condition of our study, with fixations marked with red dots. B) Smoothed fixations from panel A). The histogram of this distribution served as a reference to which the histogram of the contextualized meaning map was matched. This procedure ensures the comparability of values from both distributions by aligning these values to the same scale. C) ‘Raw’ contextualized meaning map for the scene from panel A). Since this map has not been subjects to histogram matching, color values are not comparable to values on the remaining panels. D) The map from panel C), after histogram matching but without including center bias. Interestingly, contextualized meaning maps were better predictors of fixations when they did not include the center bias (see* Soundness check 1: general predictive power of contextualized meaning maps *section). E) The map from panel C), after application of the center-bias model and subsequent subjection to histogram matching. Such maps were used in all our analyses (unless otherwise stated) because we aimed to follow the original procedure*.

#### Creating contextualized meaning maps – histogram matching

For each image, we matched the histogram of its contextualized meaning map to the histogram of the distribution obtained by smoothing human fixations registered on this image. This was done using the *imhistmatch* Matlab function. Histogram matching – also used in the original meaning maps studies – ensures that values from both distributions are directly comparable because they have been aligned to the same scale (see Fig. 2B, C, D). Similarly, as in the original study by Peacock et al. (2019), this operation was conducted after including the center-bias model in the maps.

#### Data analysis software

Data from this study was handled using Matlab R2020a (Mathworks Inc., Natick, MA) and R (R Core Team, 2020). In particular, we relied on the R packages belonging to the tidyverse collection (Wickham et al., 2019), as well as on packages jmv (The jamovi project, 2020; for running ANOVAs) and ggExtra (Attali & Baker, 2019; for generating density plots presented on Figures 5 and 6). Other R packages we used are cited in the relevant places in the text.

**Fig. 3.**
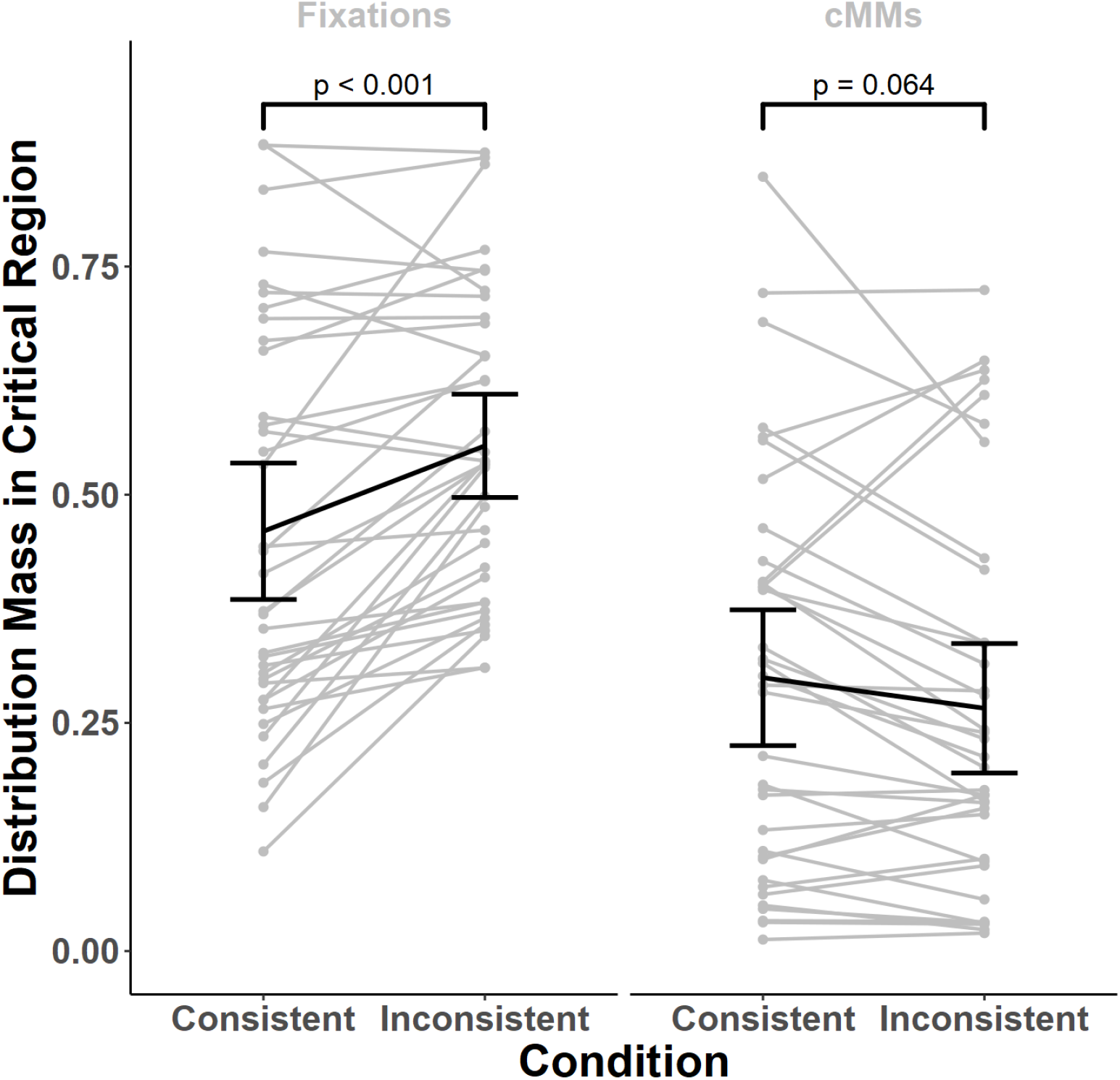
Comparison of eye-movement data and contextualized meaning maps. In each condition and for each scene, we calculated the amount of distribution-mass falling within the Critical Region (the region, in which the manipulated objects were located) divided by the Region’s area. This calculation was performed separately for smoothed fixations and contextualized meaning maps. Comparing these values between conditions revealed that observers tend to fixate the Critical Regions more when they contained semantic inconsistencies (Inconsistent condition), as compared to the situation when they did not (Consistent condition; left plot). Contextualized meaning maps (right plot, labeled cMMs) did not show this effect, as they did not attribute more meaning to semantic inconsistencies. In fact, they attributed numerically less meaning on average but this effect was not significant in a statistical sense (but see Experiment 2). Each gray line indicates a single scene, black oblique lines connect the means, black vertical lines indicate 95% confidence intervals. p-values shown on the plot were obtained using paired Wilcoxon tests, Bonferroni corrected for two comparisons.

**Fig. 4.**
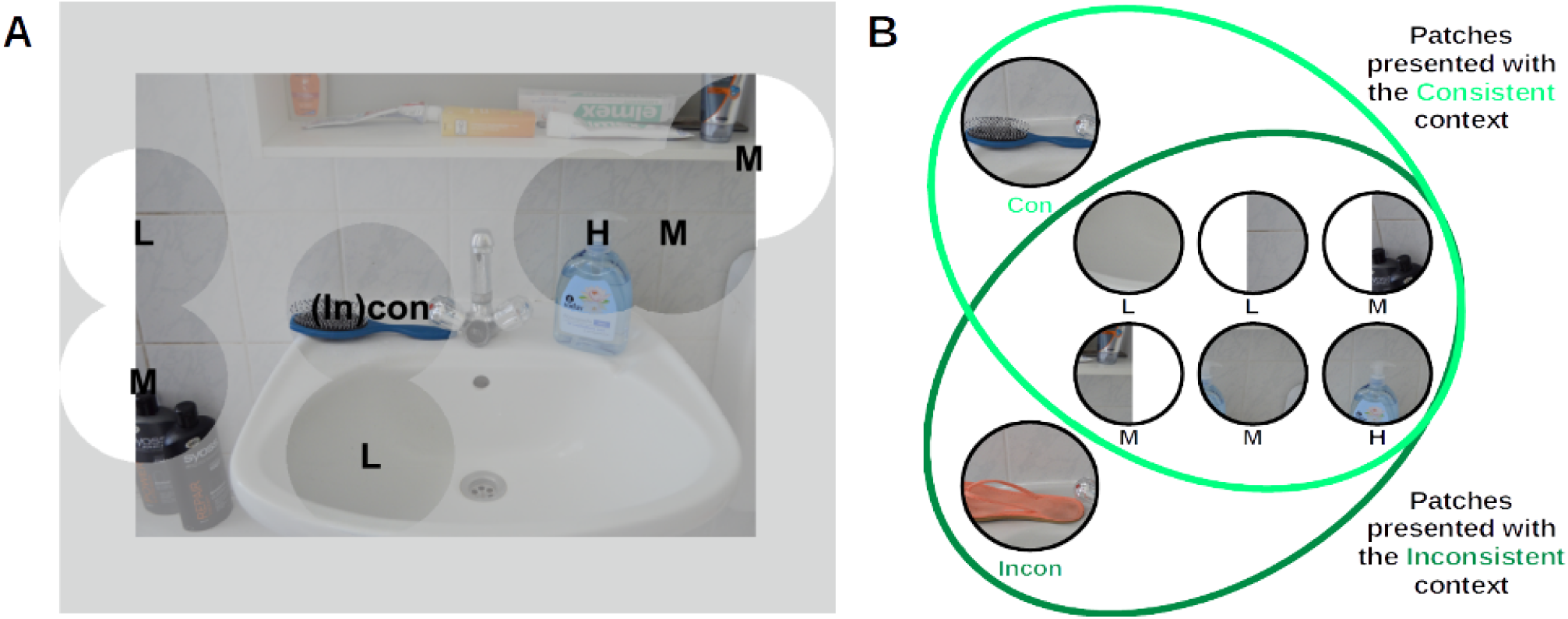
Stimulus generation for Experiment 2. A, B) In the second experiment, we tested whether patches depicting semantically inconsistent objects tend to be rated as less meaningful than their counterparts depicting consistent objects. For each scene, we selected two patches containing the consistent (Con) or the inconsistent (Incon) object. To mimic the context of the task used to generate contextualized meaning maps, we additionally included six patches that did not differ between photographs with consistent and inconsistent objects. These patches were chosen based on ratings they received in Experiment 1: on average, they had been rated as either low in meaning (labeled L on the figure, two patches), high (H, one patch) or midway between these extremes (M, three patches). Some of the patches that were selected were close to image edges and were therefore clipped. Similar to Experiment 1, each patch was presented next to either a consistent or inconsistent context scene (see panel B).

**Fig. 5.**
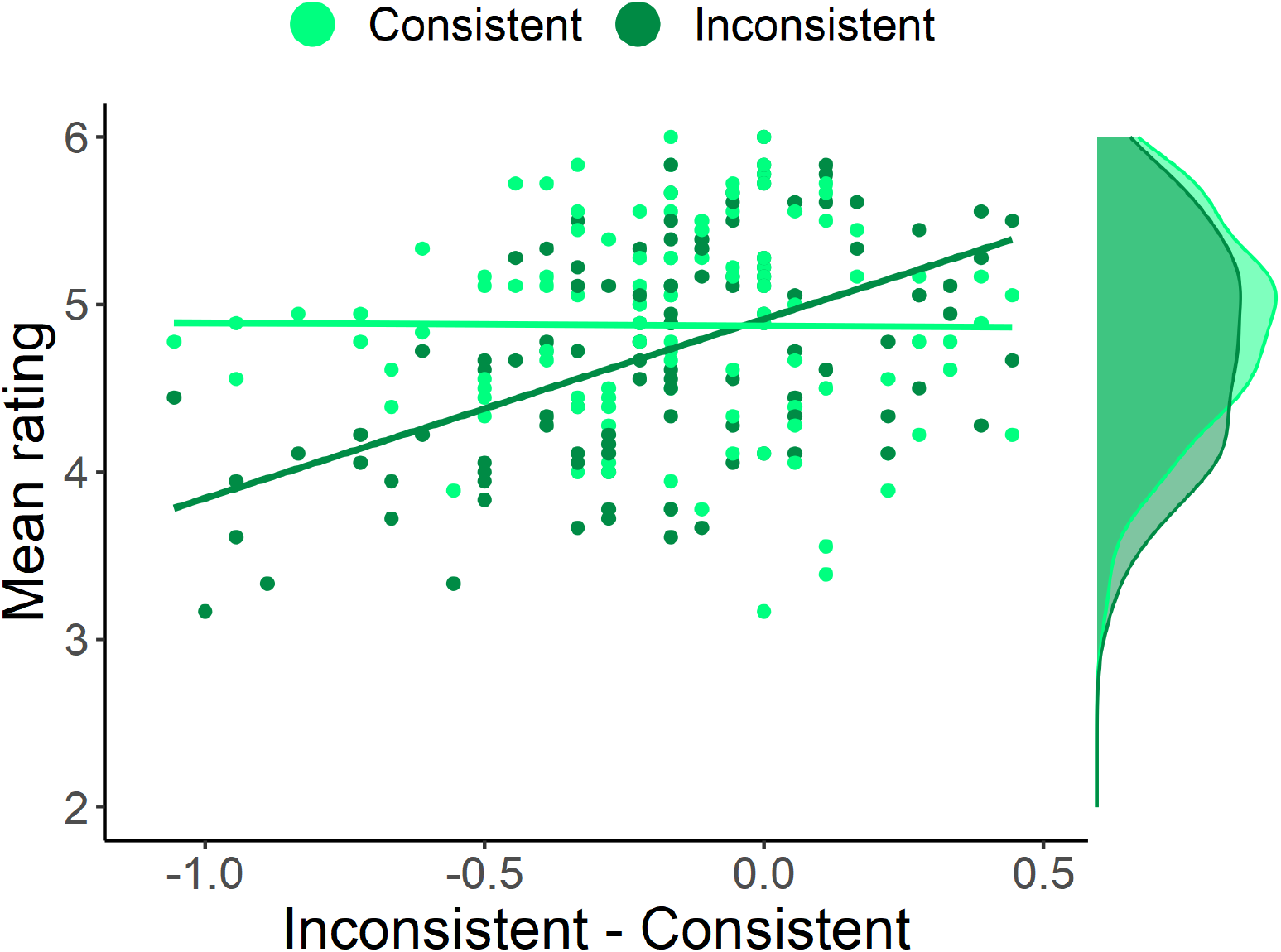
Meaningfulness ratings obtained for Con- and Incon-patches. For each rater, we averaged ratings provided for Con-patches (light-green points) and for Incon-patches (dark-green points). Next, we subtracted the average ratings for Incon-patches from Con-patches and ordered the raters according to these difference scores. The ratings for Incon-patches, but not for Con-patches, increase along this axis. Correlation analyses conducted for both types patches separately confirmed this impression: the relationship between Con/Incon differences and ratings was significant for the Incon-patches, but not for Con. Please note that this figure was generated using data not containing points identified as outliers based on their Cook’s distance (for details see main text).

**Fig. 6.**
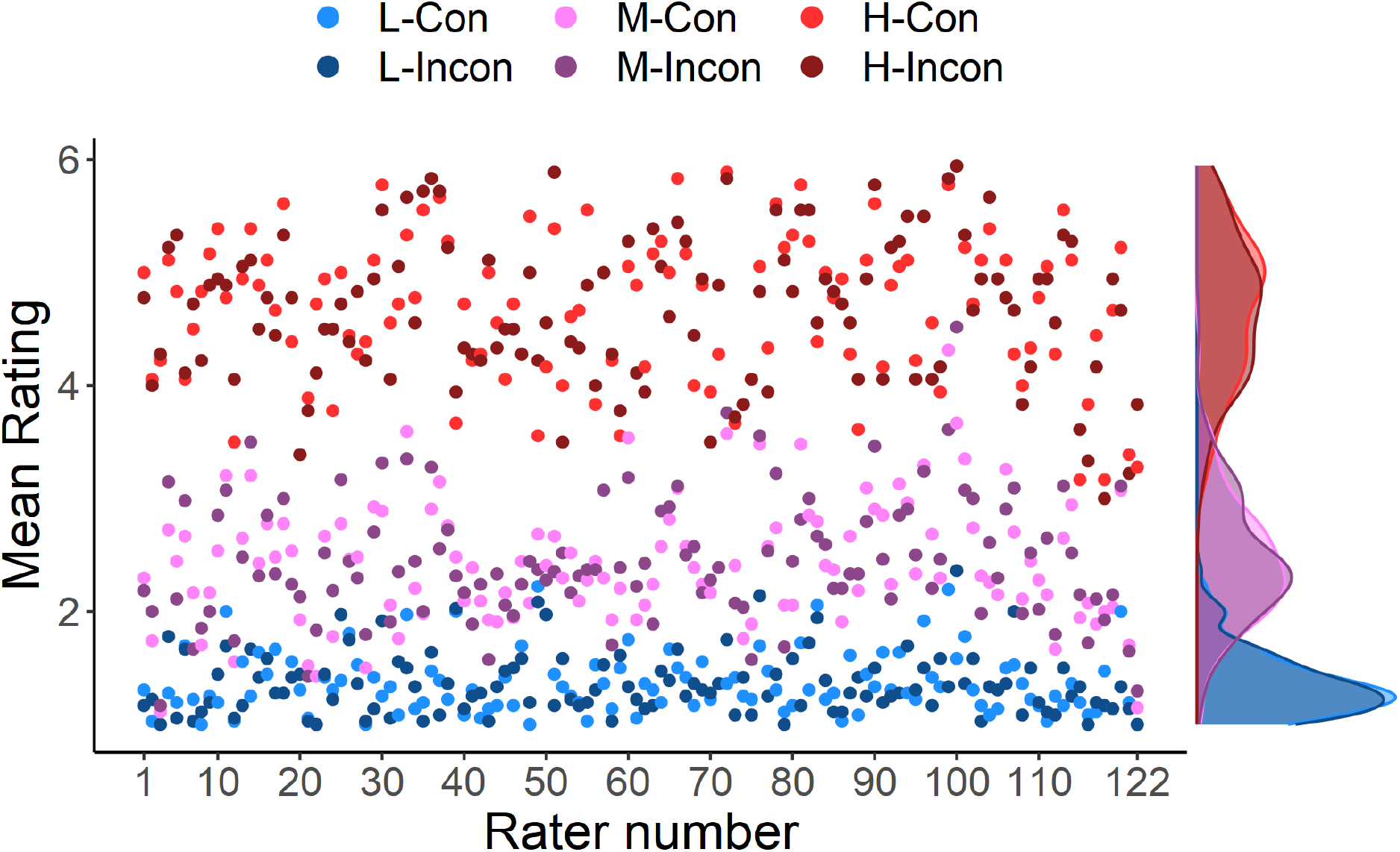
Meaningfulness ratings obtained for L-, M-, and H-patches, averaged per rater over scenes and segregated by condition. Brighter colors indicate mean ratings from the Consistent condition, darker from the Inconsistent. On the right-hand side, density plots are shown. Our analyses revealed a statistically significant main effect of patch type (L, M, and H), but no effect of condition or an interaction between condition and patch type.

#### Data and code availability

The eye movement data used in this study are openly accessible via the following link: https://zenodo.org/record/3490434). SCEGRAM stimuli are available under the following link: https://www.scenegrammarlab.com/research/scegram-database. We also share all patch-rating data and scripts for reproducing the results reported in this paper, as well as scripts and instructions for creating contextualized meaning maps (links to be provided upon publication).

### Experiment 1 – Results

#### Soundness check 1: general predictive power of contextualized meaning maps

As a soundness check, we tested how well contextualized meaning maps predicted human fixations for our stimuli: we expected them to perform at least as well as in the original study (Peacock et al., 2019). To quantify their predictive power, we applied a standard technique (Bylinskii et al., 2019), used also by Peacock and colleagues (Peacock et al., 2019): for each image, we calculated the correlation between its contextualized meaning map and smoothed fixations registered on this image. For images from the Consistent condition, the average per-image correlation was 0.60 (SD = 0.17). The average percent of the explained variance in the eye-movement data amounted to 39%. In the Inconsistent condition, contextualized meaning maps performed slightly worse (M = 0.57, SD = 0.20, 37% of the variance explained). Additionally, we investigated the effects of removing center bias from contextualized meaning maps and, interestingly, found that they performed better without it (Consistent: M = 0.71, SD = 0.13, 52% of the variance explained; Inconsistent: M = 0.66, SD = 0.17, 47% of the variance explained).

Overall, these results are similar to what is reported in the original study (Peacock et al., 2019), where the maps explained 40% of the variance in human data when center bias was included. This finding thus provides an important soundness check for our study. A lower quality of predictions in our study than in the original contextualized meaning maps study (Peacock et al., 2019) could have indicated that either the procedure of creating contextualized meaning maps is sensitive to aspects of the design which were different between our study and the original study (such as absolute image size), or that there were some technical problems with our implementation.

#### Soundness check 2: comparing contextualized meaning maps to context-free meaning maps

In our previous study (Pedziwiatr et al., 2021a), we generated original, context-free meaning maps (Henderson & Hayes, 2017) for the scenes use in the Consistent condition in the current study. As a second soundness check, we compared these original maps to the contextualized meaning maps (note that this comparison was conducted on the maps without the center bias). The average per-scene correlation between the two types of maps for the Consistent condition was M = 0.76 (SD = 0.12). Regarding the ability to predict gaze patterns, the average correlation with smoothed human-fixations was slightly higher for the context-free maps (M = 0.74, SD = 0.14 vs. M = 0.71, SD = 0.13; mean difference M = 0.03, SD = 0.01). The study that introduced contextualized meaning maps (Peacock et al., 2019) also found that contextualized and context-free meaning maps performed similarly in predicting fixations. Replicating this finding provides another soundness check for our study.

Note that the exact parameter values determining the grids used to segment images into patches differed slightly between the two types of meaning maps from our two studies. The reason for this difference is that the reports introducing the original (Henderson & Hayes, 2017) and contextualized (Peacock et al., 2019) meaning maps – on which we based our previous (Pedziwiatr et al., 2021a) and present studies, respectively – differ with respect to the reported sizes of images viewed by observers in the eye-tracking experiments (33 × 25 vs. 26.5 × 20 degrees of visual angle), yet use identical numbers of coarse and fine patches per image.

#### Sensitivity of contextualized meaning maps and eye movements to semantic manipulations

In our first main analysis, we compared contextualized meaning maps and smoothed human-fixations with respect to their sensitivity to semantic manipulations. We focused on Critical Regions – image regions which, depending on the condition, contained a semantically consistent or inconsistent objects (see *Stimuli* section for details). For each scene, we first performed histogram matching (see previous section) and then calculated the mass of each distribution (contextualized meaning maps and smoothed fixations) falling within the Critical Region and divided that value by the Region’s area for normalization. These values were then analyzed using a mixed 2×2 ANOVA with the condition (Consistent vs. Inconsistent) as a within-subjects factor and the distribution source (contextualized meaning maps vs. smoothed fixations) as a between-subjects factor (see Fig. 3). Please note that here a ‘subject’ indicates a single scene. Such an approach is typical for studies comparing fixation-prediction methods and is grounded in the observation that different observers agree to a large extent in their selection of fixation targets in images (Kümmerer et al., 2015; Wilming et al., 2011).

This analysis revealed that both the distribution sources and conditions differed from each other statistically (distribution source: F(1, 70) = 23.05, p < 0.001, ω^2^ = 0.22; condition: F(1, 70) = 5.34, p = 0.024, ω^2^ = 0.003). Importantly, however, these main effects were qualified by an interaction (F(1, 70) = 23.83, p < 0.001, ω^2^ = 0.02). For post-hoc tests, we relied on non-parametric paired Wilcoxon tests (as it is robust to the violations of the assumptions of parametric tests we observed in the data), Bonferroni-corrected for two comparisons. These tests showed that human eye-movements were sensitive to the change in semantic relationship between object and scene, as indicated by the fact that more mass of the smoothed-fixations distribution fell within the Critical regions in the Inconsistent condition compared to the Consistent condition (Inconsistent - Consistent: M = 0.09, SD = 0.12, p < 0.001). The same comparison, however, did not yield statistically significant differences for the contextualized meaning maps (M = −0.03, SD = 0.10, p = 0.064). The hypothesis that semantically-inconsistent regions carry more meaning was thus not supported by our data. Indeed, the mean rating difference, though not significant due to the correction, was in the opposite direction (consistent with the subsequent analyses and results we report below).

Further analyses yielded unexpected findings. Recall that creating contextualized meaning maps involved averaging the maps derived from coarse and fine patches. We repeated our mixed ANOVA analysis separately for each of these maps. In both cases, the pattern of results was similar to that reported in the previous section (fine maps: distribution source: F(1, 70) = 32.64, p < 0.001, ω^2^ = 0.26, condition: F(1, 70) = 0.08, p = 0.777, interaction: F(1, 70) = 31.56, p < 0.001, ω^2^ = 0.04; coarse maps: distribution source: F(1, 70) = 41.85, p < 0.001, ω^2^ = 0.30; condition: F(1, 70) = 3.71, p = 0.058; interaction: F(1, 70) = 5.87, p = 0.018, ω^2^ = 0.01). In the post-hoc tests, we did not find a difference between conditions for coarse maps (Inconsistent - Consistent: M = −0.01, SD = 0.23, p = 0.625 uncorrected). Importantly, however, we obtained an unexpected outcome in the post-hoc tests for the fine maps: these maps attributed *less* meaning to Critical Regions in the Inconsistent condition than the Consistent condition (Inconsistent - Consistent: M = −0.08, SD = 0.15, p < 0.001). Therefore, the numerical (but not statistically significant) pattern observed at the level of full maps was most likely driven by the fine maps component.

Note that these results were obtained using our custom-written, openly available implementation of meaning maps (see *Data and code availability* section). To ensure that the patterns we report above are not contingent on the specifics of our implementation, we generated contextualized meaning maps using the code shared by the authors of the original meaning maps and repeated our analyses with these maps. This code is available here: https://osf.io/654uh (*build_meaning_map* function, version uploaded to the repository on 2020/01/18). The results showed a similar pattern: both the contextualized meaning maps and their fine/coarse components attributed less meaning to the inconsistent objects (mean of the differences for full maps: M = −0.10, SD = 0.40; coarse maps: M = −0.07, SD = 0.56; fine maps: M = −0.14, SD = 0.46; note that these values are not comparable to values reported in previous analyses because here we used raw values from the *build_meaning_map* function). None of these comparisons were statistically significant (full maps: p = 0.304; coarse maps: p = 0.959; fine maps: p = 0.082), but for the fine maps this was because of Bonferroni correction for two comparisons we applied (to remain consistent with the previous analyses). Together, this analysis demonstrates that for both implementations, contextualized meaning maps do not assign more meaning to semantically-inconsistent than consistent objects.

To summarize, human eye movements changed in response to local alterations in semantic information: inconsistent objects attracted more fixations than consistent ones, and were fixated earlier. Contextualized meaning maps and their coarse component did not show this dependence on semantic information. Finally, fine maps ascribed *less* meaning to scene regions when they contained inconsistent objects, which contradicts predictions from the meaning map approach.

#### Sensitivity of patch ratings to semantic manipulations

Transforming patch ratings into contextualized meaning maps involves a number of steps, including non-linear transformations. These steps could potentially mask real, or introduce spurious between-condition differences, and for this reason, we conducted two analyses on the raw rating data. In the first analysis, we selected all patches that had an overlap of at least one pixel with the Critical Regions, and discarded the remaining patches. The ratings for patches from each condition were averaged for each scene, separately for coarse and fine patches. Averaging allowed us to account for between-scene differences in the number of patches overlapping with Critical Regions and guaranteed that the data from each scene had an equal contribution to the subsequent analyses. A comparison of these average ratings between conditions provided no evidence to suggest that between-condition differences were present in the raw data but were masked in the processes of assembling contextualized meaning maps (see Table 1 rows 1 and 4).

**Table 1:**
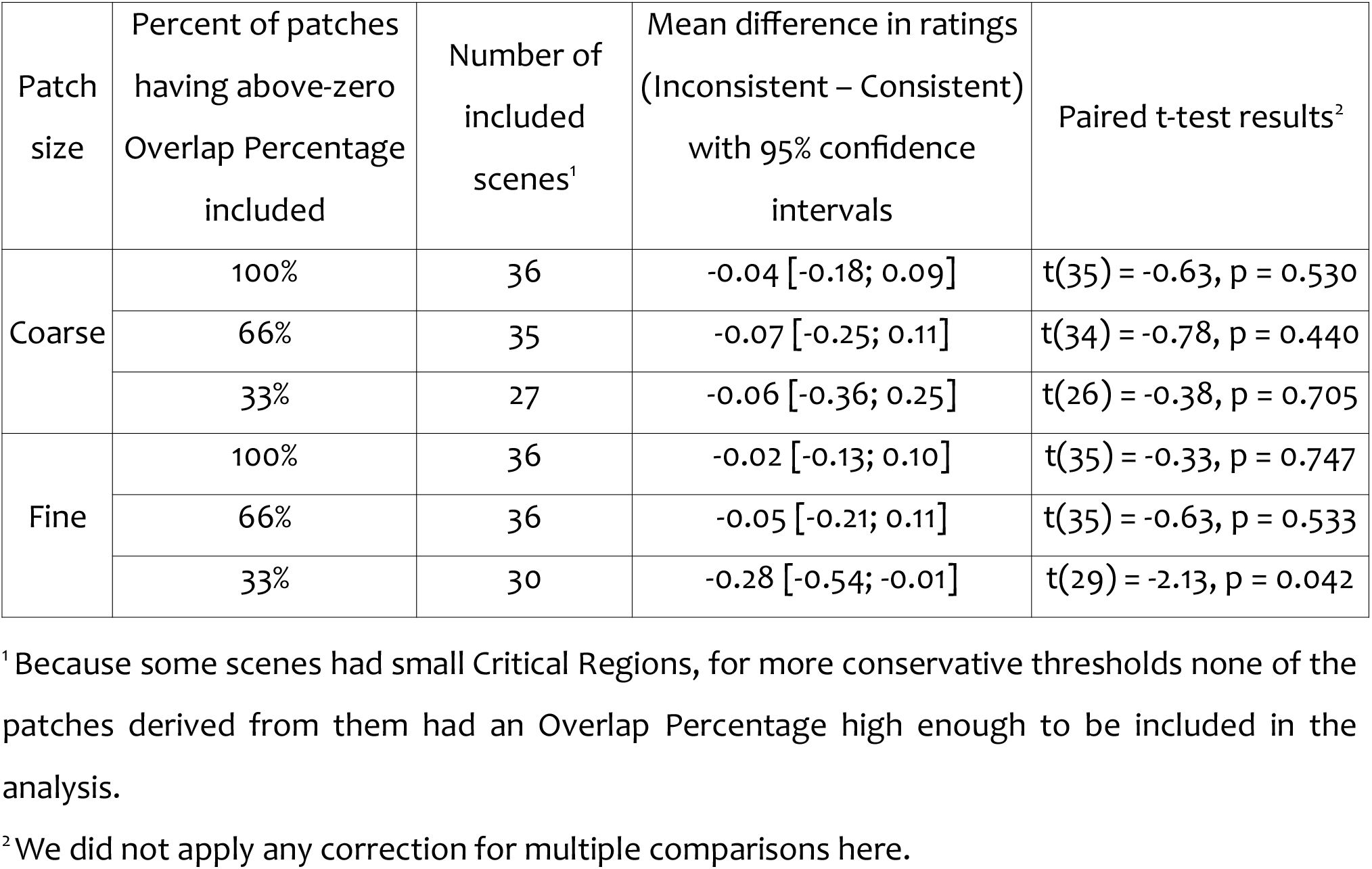
Comparison of patch ratings between conditions – statistical results

Because the above analysis included patches with at least one pixel overlap with the bounding boxes of objects, many of these patches showed only small parts of the manipulated objects, or none at all. We therefore repeated this analysis with more stringent criteria for patch inclusion. In order for a given patch to be included in this second analysis, the percentage of its area overlapping with a Critical Region (dubbed Overlap Size henceforth) had to be above a certain threshold (see Table 1). For patches of each size, we tested two threshold values. These values were selected as 34th and 67th percentiles of all above-zero Overlap Size values. For the first threshold, these values corresponded to 7% or more pixels of a patch overlapping with a Critical Region for the coarse patches, and 18% for the fine patches. Similarly, the second threshold corresponded to 21% and 56% or more overlapping pixels for coarse and fine patches, respectively. The motivation for using percentiles to determine the thresholds was to make sure that the consecutive analyses differ from each other by approximately the same percentage of retained patches: while in the first analysis we included 100% of patches which had above-zero Overlap Percentage, the thresholds resulted in including 66% (for 34^th^ percentile) and 33% (for 67^th^ percentile) of them. For each threshold and each scene, we averaged ratings of the retained patches, separately for each combination of experimental condition and patch size, and compared these per-scene values between conditions (see Table 1 for full results). Only one of the resulting tests reached statistical significance: for the most conservative threshold (i.e., with highest Overlap Size), fine patches from the Inconsistent condition were rated as *less* meaningful than their equivalents from the Consistent one. The magnitude of this difference was small: it amounted to 0.28 points on a scale from 1 to 6. The remaining five comparisons exhibited the same directionality.

#### Secondary analysis: prioritization of semantically inconsistent objects for fixation

As a secondary point of interest, we examined the temporal evolution of the influences of semantic inconsistencies on eye-movements. Other studies on the role of object-scene consistency in eye movement control yielded conflicting findings regarding whether inconsistent objects are fixated earlier or not (see Wu et al., 2014 for summary). In order to help clarifying this issue, we compared, across experimental conditions, the number of fixations required before the first fixations landed within the Critical Regions. On average, observers took 5.03 fixations (SD = 4.7) to look at the inconsistent objects for the first time, and 5.97 (SD = 5.55) for consistent (data pooled over scenes and observers). A paired Wilcoxon test indicated that this difference was statistically significant (p < 0.001). The finding that the inconsistent objects are not fixated immediately after image onset but still earlier than consistent replicates the results of a recent study by Coco, Nuthmann and Dimigen (2020). These authors supplemented gaze recordings with electroencephalography (EEG) and concluded that object semantics can be at least partially accessed via peripheral vision.

#### Summary of Experiment 1

In our first experiment, we evaluated the extent to which contextualized meaning maps and human eye-movements are sensitive to manipulations of the semantic relationship between objects and scenes. Consistent with past literature, human observers looked more at objects that are semantically inconsistent with the scene context compared to consistent objects. Contrary to predictions of the meaning map approach, however, our results provided no evidence that contextualized meaning maps assign more meaning to inconsistent than consistent objects. This insensitivity to manipulations of semantic object-scene relationships was already present at the level of the raw rating data, indicating it is not an artifact of the map generation procedure.

When we analyzed only the contextualized meaning maps resulting from ratings on fine patches, the maps assigned *less* ‘meaning’ to the Critical Region for inconsistent than consistent objects; a similar effect was observed in the raw patch data. If robust, this result would contrast with the explanation of the semantic inconsistency effect on eye movements proposed by the meaning map approach. Given that the evidence from our first experiment was based on a post-hoc subset analysis, we conducted a second experiment.

We considered two hypotheses for why we found statistically lower meaningfulness ratings for inconsistent regions in only a subset of fine patches. Firstly, it could simply be a false positive. Secondly, there might be a general but subtle tendency to rate semantic inconsistencies as less meaningful, but the subtlety of this effect might have meant that it could not be detected in ratings of coarse patches because of their low number (there were approximately 2.5 times more fine as coarse patches). The goal of Experiment 2 was to adjudicate between these two hypotheses. We created a single, well-controlled set of coarse patches derived from scenes with consistent and inconsistent objects, and collected ratings from a substantially larger sample of raters. If the reason we were unable to uncover the tendency to rate semantic inconsistencies as less meaningful in the coarse patches was due to the low number of ratings for these patches in Experiment 1, increasing the number of ratings in Experiment 2 should allow us to find this effect even in coarse patches.

### Experiment 2

#### Stimuli and design

In this experiment, we used the same 72 photographs (of 36 scenes) as in Experiment 1. For each scene, we manually selected two coarse patches that fully contained the consistent and inconsistent objects (see Fig. 4). The locations of these patches were the same in both conditions but their content changed. These patches were dubbed Con and Incon. Con-patches were derived from scenes in the Consistent condition, Incon in the Inconsistent condition. We were primarily interested in the ratings associated with these two types of patches.

To mimic the variety of patches in the rating task used for creating contextualized meaning maps and ensure that raters could use all values from the meaningfulness scale, we used the ratings from Experiment 1 to select six additional patches from each scene (see Fig. 4): two patches, which on average received the lowest meaningfulness ratings (dubbed L), one which received the highest (dubbed H), and three patches for which the ratings were midway between these extremes (dubbed M). This selection was carried out as follows. For each scene, we considered all the coarse patches that had no overlap with the Critical Region. For each location occupied by these patches, we averaged ratings across the Consistent and Inconsistent conditions. We sorted the patches according to these average ratings in an increasing order and selected two from the bottom (L), one from the top (H), and the three closest to the median (M). Therefore, we selected eight patches for each scene in total: six patches which were identical between conditions with respect to content (L, M, and H), and two patches which differed (Con and Incon). Since we expected Con- and Incon-patches to be rated as highly meaningful because they contain objects, we included only one H-patch but two L-patches in order to encourage raters to use the different scale levels with approximately equal frequency.

For stimulus presentation, each L-, M-, and H-patch was paired with the full images from both conditions. In contrast, Con- and Incon-patches were paired only with either the consistent or the inconsistent scenes, respectively. This resulted in a set of 504 patch-contexts pairs (36 scenes × 2 conditions × 6 L/M/H-patches + 36 Con-patches + 36 Incon-patches). We split this set into two equally large subsets, each containing half of the patch-context pairs from one condition and half from the other in order to avoid the situation that raters would be exposed to the same scene in both conditions. Each rater would see one of the two subsets, and thus provide ratings for 252 patches.

#### Sample-size justification

For Experiment 2, we recruited 140 raters. This sample size was largely based on the amount of resources we deemed reasonable for running this experiment. We planned to compare ratings for Con- and Incon-patches for each rater as a paired comparison (after averaging over patches; see below). After excluding 18 raters (see the *Rater inclusion criteria and inter-rater agreement* section), the resulting sample-size of 122 raters allowed detecting effects having the magnitude of Cohen’s D_z_ = 0.33 with 95% power, when using paired, two-tailed t-test and when adopting a significance level of 0.05 (as indicated by the G-Power 3.1 software; Erdfelder et al., 2009).

#### Collecting meaningfulness ratings

Data collection was conducted identically to Experiment 1. We used the same patch-rating task (with the order of stimulus presentation randomized individually for each rater) and the same method of recruiting raters (Prolific platform). The task completion times had a median of 16.12 minutes (interquartile range: 9.6).

#### Rater inclusion criteria and inter-rater agreement

We assumed that raters who followed the task instructions would agree in their ratings to a large degree. For example, we assumed that they would consistently rate M-patches higher than L-patches. Following that logic, we excluded raters whose ratings vastly disagreed with the ratings provided by the majority of participants. We operationalized this idea by first measuring the agreement of ratings within each possible pair of raters who had viewed the same subset of patches using Krippendorff’s α (A. F. Hayes & Krippendorff, 2007; Krippendorff, 1970). Values of α span from negative values to 1, where 1 indicates perfect agreement, 0 indicates the degree of agreement achievable by chance, and negative values indicate systematic disagreement. We calculated pairwise α for our raters using the function kripp.alpha from the R package irr (Gamer et al., 2019), with the option *scaleType* set to ‘interval’ (setting it to ‘ordinal’ did not influence the pattern of results). Next, for each rater, we averaged the α values from all pairs to which this rater belonged. These per-rater average α values (dubbed R_α_ henceforth) indicated the degree to which a given rater agreed with other raters who rated the same subset of patches. We visually inspected the histogram of R_α_ values calculated for all raters and decided that in our final sample, we would include only raters having R_α_ larger than 0.40. This resulted in excluding 18 raters and retaining 122 (importantly, our main results do not depend on this step – see *Influence of data exclusions* section). The average R_α_ for the retained raters was 0.70 (SD = 0.06). Additionally, we calculated R_α_ values for the excluded raters only. These values indicated the agreement being close to the chance level (mean = −0.06, SD = 0.20) which means that these raters were most likely responding at random, rather than using a common rating strategy, consistently differentiating them from the majority of our sample.

### Experiment 2 – Results

#### Patches that were manipulated between conditions (Con and Incon)

The main focus of Experiment 2 was to assess whether objects that are semantically inconsistent with the scene context are rated differently with respect to the amount of meaning they convey compared to consistent objects. Recall that each rater saw both Con- and Incon-patches, but not the same scene in both conditions. We averaged ratings over patches in each condition to yield a Con- and Incon-average rating for each rater, then compared them with a paired-samples t-test. In line with the preliminary findings of Experiment 1, the results demonstrate that semantically inconsistent objects were rated as less meaningful compared to consistent objects. The absolute magnitude of this effect was small (mean of the differences: M = −0.21, 95% CI [−0.14, −0.28]; median: −0.17) but statistically significant (t(121) = 5.80, p < 0.001).

To assess the contribution of the consistent vs. the inconsistent condition to this effect in a subject-by-subject approach, we ordered the raters by the difference between their average rating for Con- and Incon-patches. As shown in Fig. 5, this difference seems to be largely due to changes in ratings of inconsistent patches: while there was no clear subject-by-subject difference in the ratings for Con-patches, raters who contributed to the group-level effect showed decreased ratings for D-Incon patches. This impression was corroborated by a statistical analyses that showed a significant correlation between Con/Incon differences and the Incon ratings (r(111) = 0.52, 95% CI [0.37; 0.64], p < 0.001), but no such relationship for Con ratings (r(111) = −0.01, 95% CI [−0.19; 0.18], p = 0.928). Note that – for each analysis separately – we excluded points which had a Cook’s distance higher than 3 times the mean Cook distance for all points. For Con ratings, this exclusion threshold amounted to 0.02 (0.03 for Incon) and resulted in 9 exclusions (also 9 for Incon). We applied these exclusion criteria because the initial inspection of the data suggested that, in each case, the effects might be driven by a small number of points, which would have a disproportionately large influence on regression. However, repeating the analyses with all the data included resulted in the same pattern of outcomes (Incon: r(120) = 0.50, 95% CI [0.36; 0.62], p < .001; Con: r(120) = −0.08, 95% CI [−0.25; 0.10], p = 0.398).

These findings suggest that there is high consistency across raters regarding their evaluation of the meaningfulness of objects that are semantically consistent with their scene context. Ratings for inconsistent objects, in contrast, revealed considerable variability in rater behavior. Different individuals tended to rate these objects as either lower, similar, or higher in meaning than the consistent objects. Ultimately, this difference not only offers interesting insights into individual differences but also suggests that the group-level effect is mainly driven by changes in the ratings of inconsistent objects.

Our final analysis focused on individual scenes, rather than individual raters, comparing ratings for Con- and Incon-patches derived from the same scenes. For each scene, we conducted a separate between-subjects Welsh test comparing ratings received by Con- and Incon- patches, similar to the analysis conducted for L/M/H-patches. Without correction for multiple comparisons, 13 out of 36 of these tests yielded statistically significant results (this number was reduced to 3 after applying the correction). Out of these 13 cases, in 12 (33% of all scenes) the Incon-patch was rated as less meaningful than the Con-patch. These findings suggest that the tendency of Incon-patches to be rated as less meaningful than Con-patches was observable at the level of scenes too, which corroborates the finding from the rater-level analysis.

In summary, our main analyses demonstrate two key findings: first, we show that semantically inconsistent objects are rated as less meaningful compared to consistent objects. Second, the size of this effect shows marked individual differences between raters.

#### Influence of data exclusions

Recall that at the initial stage of our analyses we excluded 18 raters (see *Rater inclusion criteria and inter-rater agreement* section). In order to make sure that our conclusions do not critically depend on this step, we repeated all the analyses from the previous section with the data from all raters recruited for Experiment 2. This operation did not change the pattern of our results (comparison of ratings for Con- and Incon-patches: t(139) = 5.99, p < 0.001, mean of the differences M = −0.20, 95% CI [−0.13; −0.27]; correlation for Con-patches: r(138) = −0.10, 95% CI [−0.26; 0.07], p = 0.242; correlation for Incon-patches: r(138) = 0.37, 95% CI [0.21; 0.50], p < 0.001).

#### Soundness check: patches that were identical between condition (L, M, and H)

As a soundness check, we tested whether L, M, and H-patches were rated as low, medium and high in meaning, respectively. We used Page’s test, a non-parametric, rank-based statistical test assessing the ordering of values obtained in repeated measurements (Page, 1963), and compared the null hypothesis that there were no differences between ratings for all three types of patches against the alternative stating that L-patches (mean rating M = 1.36, SD = 0.28) were rated lower than M-patches (M = 2.44, SD = 0.55) which, in turn, were rated lower than H-patches (M = 4.69, SD = 0.65). We implemented the test with the R package crank (Lemon, 2019) and conducted it separately for patches from the Consistent and the Inconsistent conditions. In both cases the results were statistically significant (and identical numerically: L = 1708, p < 0.001) which indicated that the pattern of obtained results matched our expectations.

To evaluate whether the presence of consistent or inconsistent objects in a scene affect the ratings for all patches in that scene, we analyzed whether ratings for L-, M-, and H-patches differed between consistent and inconsistent conditions. For each rater, we averaged ratings provided for each of these patch types per condition (see Fig. 6), and analyzed the averages with a 2×3 repeated-measures ANOVA (with a Greenhouse-Geisser correction) with the two within-subjects factors Condition (Consistent and Inconsistent) and Patch-Type (L-, M-, and H-patches). As expected based on the preceding findings, this analysis also showed that ratings differed according to patch type, as indicated by a main effect for this factor (F(1.57, 190.25) = 2530.65, p < 0.001). The other main effect and the interaction showed no significant differences (Condition: F(1, 121) = 0.02, p = 0.883; interaction: F(1.35, 163.16) = 0.77, p = 0.418), showing that average ratings for L-, M- and H-patches did not differ depending on whether the full scene contained a consistent or inconsistent object.

In a final analysis of the L-, M-, and H-patches, we focused on potential differences between individual scenes. The previous analyses reported in this section averaged patch ratings per rater over scenes. In our final analysis, we took a different approach and compared ratings provided for individual L-, M-, and H-patches across conditions. Individual patches were rated by a separate set of raters in the Consistent and Inconsistent conditions (see section *Stimuli and design* section). We therefore used a between-subjects Welch test to compare the ratings for each patch individually across conditions and found statistically significant differences only for 2 patches (out of 216), derived from 2 different scenes. Therefore, in the vast majority of cases, the condition from which the context image was derived did not influence the ratings for individual patches.

Overall, these control analyses have two implications. First, they indicate that the raters adopted the expected rating strategy, as suggested by the expected ordering of values for L-, M-, and H-patches. Second, exchanging a single object that is semantically consistent with the scene for an inconsistent object did not have a general effect on the rating of patches that did not contain the manipulated object, neither on average nor on a scene-by-scene level.

## Discussion

Human fixations are attracted to objects that are semantically inconsistent with the scene within which they appear. One possible explanation of these effects is that these objects carry increased meaning, which causes people to look at them more. This hypothesis has gained increasing attention with the development of meaning maps, a novel tool to index the distribution of meaning across an image (Henderson & Hayes, 2017, 2018; Peacock et al., 2019). In two experiments, we tested if semantically inconsistent objects indeed carry more meaning as measured by contextualized meaning maps (Peacock et al., 2019), which have been designed to capture such contextual effects. First, we created contextualized meaning maps for images of scenes containing objects that were either semantically consistent or inconsistent, and compared these maps to eye-movement data. While observers looked more at inconsistent compared to consistent objects, contextualized meaning maps did not attribute higher amounts of meaning to the former than to the latter. In fact, we found preliminary evidence to suggest that the same scene location might be indexed as less rich in meaning when it contains semantic inconsistency. In a second experiment, we therefore asked a substantially larger number of raters to provide meaningfulness ratings for a carefully controlled set of image patches, including patches that showed semantically consistent or inconsistent objects. The results of this second experiment provide evidence suggesting that human observers have a tendency to judge objects that are semantically inconsistent with the scene as slightly less meaningful than their consistent counterparts.

The tendency of human observers to look more at semantically inconsistent objects is considered to be a prototypical example of semantic influences on eye movements. Several previous explanations of this effect implicitly or explicitly assume that semantic inconsistency increases the amount of (semantic) information, or meaning that is conveyed (Henderson, 2011; Henderson et al., 1999; Loftus & Mackworth, 1978). This interpretation has been strongly expressed within the recently developed meaning map approach (Henderson & Hayes, 2017, 2018; Peacock et al., 2019 see also Henderson et al., 2019 for review). In contrast to this notion, our direct evaluation of contextualized meaning maps suggests that, while they show a good overall ability to predict human gaze patterns, they are unable to predict influences of semantic inconsistencies, showing no difference between our Consistent and Inconsistent conditions. Therefore, contextualized meaning maps fail to capture at least one context-based semantic influence on eye-movement control.

It is important to highlight the fact that a conceptualization of meaning in terms of object-context relationships is by no means exhaustive. Other conceptualizations have been proposed (T. R. Hayes & Henderson, 2021; Hwang et al., 2011; Rose & Bex, 2020) and the idea that there might be several subtypes of meaning that are important for eye movements has been suggested by other authors (Henderson et al., 2018; Henderson & Hayes, 2018). Our findings indicate that contextualized meaning maps and patch ratings do not capture the effect of semantic object-scene relationships on eye movements, but they might measure other types of meaning (see also Henderson et al., 2021). The critical question therefore is what type of gaze-relevant meaning they might measure.

Answering this question is impeded by the fact that it is far from clear what raters are doing when asked to provide meaningfulness judgments for image patches. In both experiments, we used the instructions from the original contextualized meaning maps study by Peacock et. al (2019). These instructions define meaningfulness in rather vague terms by linking it to informativeness and recognizability. Raters are instructed as follows: *“We want you to assess how “meaningful” an image is based on how informative or recognizable you think it is*”. Our study shows that, at the group-level, such instructions lead to lower meaningfulness ratings for objects that are semantically inconsistent with the scene context. One possible explanation for this result is that raters find it more difficult to recognize inconsistent objects (“What is that on the sink there? A shoe?”), and might therefore rate the meaningfulness of the patch lower (emphasizing the *“recognizable”* component of the definition of meaningfulness used by the meaning maps approach). Also note that the ambiguity of the instruction may cause higher inter-subject variability in the inconsistent condition because raters might be unsure about how to interpret the image manipulations in the context of the instructions.

Other instructions would likely lead to qualitatively different findings. For instance, imagine observers were given identical instructions to those used in our study except that they were also told that the images in the study show crime scenes. It seems plausible that raters would pick out the semantically inconsistent objects as being particularly meaningful in this context (emphasizing the *“informative”* aspect of the instruction). Adjusting task instructions (and, potentially, the parameters of grids used for segmenting scenes into patches) systematically in a wide range of cases in order to maximize the predictive power of the resulting maps might be an interesting research direction. However, such an approach would entail treating meaning maps not as a tool to measure the distribution of semantic information in scenes, but as another method of predicting human fixations: a crowd-sourced saliency model. That is, a method which prioritizes the quality of predictions over both the interpretability of mechanisms generating these predictions (i.e. the ability to identify factors determining the accuracy of predictions) and the explanatory power (i.e. the amount of gained insight into human oculomotor control).

Alternatively, the variability in responses in the patch-ratings task in its current form makes this task a potentially interesting tool for indexing individual differences (Hedge et al., 2018). While we currently lack clarity regarding the processes underpinning the selection of rating values, further research, combining the patch-rating task with other measures, might shed more light on this issue, and thereby on individual differences in how the content of natural scenes is processed. This topic is still understudied in the context of eye movements, despite the evidence showing that such individual differences exist (De Haas et al., 2019; see also Kröger et al., 2020).

Given the limitations of human rating data, current developments in computational approaches might provide alternative methods that could contribute to a better understanding of the role of high-level factors in eye-movement control, including semantic information and meaning. A number of authors have attempted to develop indices of these high-level aspects of visual input by applying techniques to images that have originally been developed in natural-language processing (T. R. Hayes & Henderson, 2021; Hwang et al., 2011; Lüddecke et al., 2019; Rose & Bex, 2020; Treder et al., 2020), in particular in the field of distributional semantics (Harris, 1954). While these computational methods come with their own limitations, they have a number of advantages over human rating data: they are comparably inexpensive, fast, and easy to use, and can comfortably be applied to large image data sets due to their automation. Moreover, computational tools have the potential to be less opaque compared to human rating data, and might be more amenable to detailed analyses of which aspects of high-level scene content contributes to eye-movement control. For instance, the finding that humans look more and longer at semantically inconsistent objects might be based purely on a statistical analysis of object co-occurrences in visual scenes (see Wang et al., 2010). Not surprisingly, recent analyses of image datasets with more than 20 000 images indicate that different scene categories indeed show a highly consistent clustering of object types (Treder et al., 2020), and the oculomotor system might exploit these regularities for outlier detection. This interpretation of the influence of object-scene inconsistencies on eye movements is similar in spirit to earlier notions of saliency (Bruce & Tsotsos, 2009), but transfers this idea from a low-level (feature-based) to a high-level (object- and scene-based) analysis of the visual input. While – most likely – being an important contributor, co-occurrence *per se* does not necessarily amount to a semantic relationship between objects, or meaning. And some computational approaches, such as the one developed by Treder and colleagues (Treder et al., 2020), might have the potential to determine whether oculomotor control relies purely on basic co-occurrence or transforms these raw data further into a type of information that is closer to what we might label ‘meaning’.

To summarize, introducing semantic inconsistencies to a scene region by replacing a semantically consistent object with one that is semantically inconsistent did not increase the amount of meaning attributed to this region by contextualized meaning maps, despite increasing the number of human fixations landing on this region. Therefore, even though the maps predicted human fixations well for scenes containing only consistent objects, they are not able to account for semantic influences on human gaze-allocation linked to semantic object-context inconsistencies. In fact, data from both of our experiments provide evidence suggesting that human observers might have the tendency to rate semantically inconsistent objects as slightly less meaningful than their consistent counterparts. Our results further highlight the need for improved conceptualization and methods to investigate the role of semantic information in human oculomotor control.

## CRediT author statement

**M.P.**: Conceptualization, Methodology, Software, Investigation, Formal Analysis, Writing – Original Draft, Writing – Review & Editing

**M.K., T.W., M.B.**: Conceptualization, Writing – Review & Editing

**C.T.:** Conceptualization, Formal Analysis, Resources, Writing – Original Draft, Writing – Review & Editing, Supervision

## Acknowledgments

We would like to thank Antje Nuthmann and Tom Freeman for their comments on an earlier version of the manuscript. This work was supported by the German Federal Ministry of Education and Research (BMBF): Tübingen AI Center, FKZ: 01IS18039A and the Deutsche Forschungsgemeinschaft (DFG, German Research Foundation): Germany’s Excellence Strategy – EXC 2064/1 – 390727645 and SFB 1233, RobustVision: Inference Principles and Neural Mechanisms.

